# The effect of acquisition resolution on orientation decoding from V1: comparison of 3T and 7T

**DOI:** 10.1101/305417

**Authors:** Ayan Sengupta, Oliver Speck, Renat Yakupov, Martin Kanowski, Claus Tempelmann, Stefan Pollmann, Michael Hanke

**Affiliations:** Department of Experimental Psychology, Institute of Psychology, Otto-von-Guericke University, Universitätsplatz 2, 39016 Magdeburg, Germany; Sir Peter Mansfield Imaging Centre, University of Nottingham, UK; Department of Biomedical Magnetic Resonance, Institute for Experimental Physics, Otto-von-Guericke University, Leipziger Str. 44, 39120 Magdeburg, Germany; Center for Behavioral Brain Sciences, Otto-von-Guericke University, Universitätsplatz 2, 39016 Magdeburg, Germany; Leibniz Institute for Neurobiology, Brenneckestr. 6, 39118 Magdeburg, Germany; German Center for Neurodegenerative Disease (DZNE), site Magdeburg, Leipziger Straβe 44, Germany; Psychoinformatics lab, Institute of Psychology, Otto-von-Guericke University, Universitätsplatz 2, 39016 Magdeburg, Germany; Department of Neurology, Otto-von-Guericke University, Leipziger Str. 44, 39120 Magdeburg, Germany

**Keywords:** functional magnetic resonance imaging, acquisition resolution, decoding, field strength comparison, visual orientation, spatial filter, 3T, 7T

## Abstract

Previously published results indicate that the accuracy of decoding visual orientation from 7 Tesla fMRI data of V1 peaks at spatial acquisition resolutions that are routinely accessible with more conventional 3 Tesla scanners. This study directly compares the decoding performance between a 3 Tesla and a 7 Tesla dataset that were acquired using the same stimulation paradigm by applying an identical analysis procedure. The results indicate that decoding models built on 3 Tesla data are comparatively impaired. Moreover, we found no evidence for a strong coupling of BOLD signal change magnitude or temporal signal to noise ratio (tSNR) with decoding performance. Direct enhancement of tSNR via multiband fMRI acquisition at the same resolution did not translate into improved decoding performance. Additional voxel selection can boost 3 Tesla decoding performance to the 7 Tesla level only at a 3 mm acquisition resolution. In both datasets the BOLD signal available for orientation decoding is spatially broadband, but, consistent with the size of the BOLD point-spread-function, decoding models at 3 Tesla utilize spatially coarser image components.

## 1. Introduction

Multivariate pattern analysis, and more specifically decoding analysis, has become part of the standard toolkit of neuroimaging data analysis (Haxby, 2012). However, many applications of decoding analysis primarily aim at testing the ability to decode a particular set of conditions, for example, from activity patterns of a specific brain region, without investigating the nature of the signal that contributes to decoding. Consequently, there are still many open questions regarding optimal modeling and pre-processing of imaging data. Likely the most extensively studied paradigm on this topic is the decoding of representations of visual orientation from early visual cortex (e.g., Op de Beeck, 2010; Freeman et al., 2011; Alink et al., 2013; Freeman et al., 2013). In this context, Sengupta et al. (2017) investigated the effect of acquisition resolution, as well as type and magnitude of spatial filtering on orientation decoding from V1 using ultra-high field 7 Tesla fMRI data recorded at four different resolutions (0.8 mm, 1.4mm, 2mm and 3mm iso). Among the tested acquisition resolutions, the 2 mm data were found to yield the best decoding accuracy (even when considering several preprocessing strategies of higher resolution scans) – a fairly conventional resolution that is well within the capability range of more common 3 Tesla MR scanners. Moreover, Mandelkow et al. (2017) reports that decoding of naturalistic stimuli from 7 Tesla data doesn’t not generally benefit from a resolution increase from 2 mm to 1.2 mm, and decoding accuracy was found to peak broadly around a Gaussian-smoothed resolution of 3 mm full width at half maximum (FWHM). These findings contradict the expectation that high-resolution fMRI should lead to better decoding performance, as, for example, a substantial reduction in the partial volume effect at higher resolution leads to a larger BOLD signal magnitude (Weibull et al., 2008), and therefore should allow for an improved capture of fine-grained activitation patterns. However, Mandelkow et al. (2017) reports that decoding of naturalistic stimuli from 7 Tesla data doesn’t not generally benefit from a resolution increase from 2 mm to 1.2 mm, and decoding accuracy was found to peak broadly around a Gaussian-smoothed resolution of 3 mm full width at half maximum (FWHM).

This raises the question what impact the magnetic field strength has on the performance of decoding models (Formisano and Kriegeskorte, 2012). A simulation by Chaimow et al. (2011) predicted that the optimal decoding accuracy of, for example, ocular dominance, can be found between 2-3 mm voxel size at 3 Tesla because technical and physiological factors place the optimal acquisition resolution for decoding at a substantially coarser scale than the underlying columnar structure. Mandelkow et al. (2017) reports that decoding of naturalistic stimuli from 7 Tesla data Consequently, one could hypothesize that equally performant decoding models can be built on 3 Tesla fMRI data, compared to 7 Tesla. However, while the performance differences across resolutions at a constant field strength could be similar, the higher BOLD sensitivity at 7 Tesla may still result in generally higher decoding accuracies. This study explores this hypothesis by applying the same stimulation and analysis paradigm as in Sengupta et al. (2017) on matching fMRI data acquired at 1.4 mm, 2 mm, and 3 mm with a 3 Tesla MR scanner. In addition, the role of the temporal signal to noise ratio of fMRI time series as a predictor of decoding performance that was proposed in Sengupta et al. (2017) was explicitly tested here, and the spatial scale of the captured orientation-related signal was, again, examined through post-acquisition volumetric Gaussian filtering.

## 2. Materials and methods

### Participants

Seven healthy right-handed participants (five male) with normal or corrected to normal vision were recruited from the subject-pool of the *studyforrest* project (Hanke et al., 2014, 2015). Five of those also volunteered for the previous 7 Tesla experiment (Sengupta et al., 2017), sub-04 and sub-18 were replaced by sub-09 and sub-10 due to unavailability. Before every scanning session, participants were provided with instructions for the experiment and signed an informed consent form. The study was approved by the Ethics Committee of the Otto-von-Guericke University. Participants received monetary compensation.

### Stimulus and Experimental Design

In order to keep parity between the 7 Tesla and 3 Tesla experiments, the stimulus was kept identical to that described in Sengupta et al. (2017), except for adjustments to match the different stimulation setup. Flickering sine-wave orientated gratings (flicker frequency = 4 Hz, constant spatial frequency 1.4 cycles per degree of visual angle with 100% contrast) were displayed in both hemifields on medium gray background in form of semi-annular patches (0.8°-7.6° eccentricity, 160° width on each side with a 20° gap along the vertical meridian). Orientation gratings (0°, 45°, 90°, or 135°) were displayed with random phase (0, 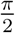, π, or 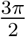 degrees) changed at a frequency of 4 Hz.

Stimulation and response logging were implemented using PsychoPy (v1.79; Peirce, 2008) on a standard PC running (Neuro)Debian (Halchenko and Hanke, 2012). The stimulus was displayed on a rear-projection screen (1140×780 pixels, 18cm wide), 60Hz video refresh rate) placed at a viewing distance of 35 cm. In order to keep the participants’ attention focused and to minimize eye-movements, they performed a center fixation task that was unrelated to the stimulation with oriented gratings. Participants were asked to fixate on a Landolt-C stimulus (radius 0.12°, left or right opening (0.048°) presented at the center of the screen. At random intervals in each run the Landolt-C was shown and the participants had to respond to the direction of the opening of the probe by pressing one of two buttons corresponding to a left or right opening. The mean accuracy for this task was 90.1% (std=9.63) correct across all participants.

Acquisition of fMRI for the different spatial resolutions was performed in three separate sessions, in random order for each participant. Each participant performed ten runs of the experiment for each spatial resolution. Each run comprised of 30 trials (8s duration; 4min total run duration). There were 3s of the flickering stimulus at the beginning of each trial followed by a 5s inter-trial interval. Details of the generation of the pseudo-random orientation stimulus sequence were identical to Sengupta et al. (2017).

### MRI data

The objective for functional data acquisition was to obtain BOLD fMRI data from early visual cortex at three different resolutions with minimal differences in acquisition parameters with respect to the previous study and across acquisition resolutions, give two a-priori constraints: 1) sufficient spatial coverage of the ROI, and 2) identical temporal sampling frequency (TR) across resolutions.

T2*-weighted echo planar images (EPI) (TR/TE=2000/30ms, FA=90°) of the occipital lobe were acquired using a 3 Tesla Siemens Prisma scanner with the 52 head coil elements of a 64 channel head neck coil (Siemens, Erlangen, Germany). Slices, oriented approximately parallel to the calcarine sulcus (on a tilted axial plane), were acquired for three different spatial resolutions, i.e. 3 mm isotropic (FoV=216mm, matrix size 72×72, 36 slices, GRAPPA accel. factor 2), 2mm isotropic (FoV=216mm, matrix size 108×108, 32 slices, GRAPPA accel. factor 2) and 1.4mm isotropic (FoV=210mm, matrix size 150×150, 28 slices, GRAPPA accel. factor 3). All EPI scans implemented ascending slice acquisition order and used a 10% inter-slice gap to minimize cross-slice excitation. All acquisition sequences used anterior-posterior phase encoding. For each experiment run, 120 volumes were acquired and 10 separate scans (one for each experimental run) were performed for each subject.

For the purpose of evaluating the impact of the temporal signal-to-noise-ratio (tSNR) on decoding accuracy, an additional acquisition was performed at 2 mm resolution. While all other acquisitions employed parallel imaging (via GRAPPA; Griswold et al., 2002) to reduce acquisition time at the cost of decreased SNR (for a parallel imaging acceleration factor of *R*, SNR is expected to be reduced by at least a factor of 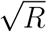; Glockner et al., 2005), this acquisition was performed without GRAPPA acceleration, but instead with simultaneous multi-slice or multiband EPI (Moeller et al., 2010) in order to maintain the scan timing for the same field of view (SMS=2, TR=2000 ms, FA=90°, 7/8 k-space, 120 volumes).

For comparison with fMRI data acquired at a different magnetic field strength, the 7 Tesla data from Sengupta et al. (2017) with spatial resolutions 1.4 mm, 2.0 mm and 3. mm isotropic voxel size were re-used in this study. The dataset is publicly available from OpenfMRI (accession: ds000113c).

Structural images for all participants were previously acquired as a part of the *studyforrest* project and were re-used in this study. This includes the reconstruction of the individual cortical surface mesh models (Hanke et al., 2014).

### Region of interest localization

All participants who volunteered for this study previously underwent retinotopic mapping (Sengupta et al., 2016, data and results publicly available from OpenfMRI ds000113d). Using these data, V1 ROIs were manually delineated on the inflated cortical surface for each participant, and were then converted to volumetric masks for each acquisition resolution. For the participants of the previous 7 Tesla experiment, the exact same ROI definitions were used. For the two new participants, ROIs were defined using the identical procedure.

### Data analysis

Data preprocessing and analysis implementation were largely identical to those reported in Sengupta et al. (2017). However, while motion-correction of the 7 Tesla data was performed as part of a scanner-side distortion correction routine (In and Speck, 2012), the 3 Tesla data were motion corrected using MCFLIRT in FSL (v5.0.8; Smith et al., 2004).

Based on our previous findings that the specific nature of spatial filtering strategies (volumetric vs. surface-based, pre-filter vs. post-filter masking) has minimal effect on the results (Sengupta et al., 2017), we decided to exclusively use the strategy of applying spatial filters to an entire EPI volume, and subsequently analyze the data from voxels corresponding to the ROI (analog to, e.g., Swisher et al., 2010).

In order to test the spatial scale of orientation-related signal, a variety of spatial filters were applied to the data for repeated analysis. As in Sengupta et al. (2017) the magnitude of spatial filtering is reported as the size of the Gaussian filter kernel described by its FWHM size in millimeters. The implementation of spatial filtering was identical to the previous study. *Low-pass* (LP) 3D Gaussian spatial filtering was performed with a kernel size of up to 20 mm FWHM in steps of 1 mm. Complementary *high-pass* (HP) filtered images were computed by subtracting the respective LP filtered image of a particular filter size from the unfiltered image. These images include only the fine-scale orientation specific fMRI patterns after removal of the low-frequency components. *Band-pass* (BP) filtered images were computed by a difference-of-Gaussians filtering approach (Alink et al., 2013) by subtracting LP filtered images for two filter sizes (1mm distance) from each other. Corresponding *band-stop* (BS) images where computed by subtracting the BP image from the original unfiltered image.

Decoding analysis was performed with PyMVPA (v2.6.3; Hanke et al., 2009) on a compute cluster running (Neuro)Debian (v9.3; Halchenko and Hanke, 2012). For feature extraction, a general linear model (GLM) was fitted to the BOLD fMRI time series from an individual experimental run using the GLM implementation in NiPy (v0.4.1; Millman and Brett, 2007) while accounting for serial correlation with an autoregressive term (AR1). As in the previous study, the GLM design matrix included hemodynamic response regressors for each stimulus condition, separately for both hemifields, and corresponding temporal derivatives, six nuisance regressors for motion (translation and rotation), and polynomial regressors (up to 2nd-order) modeling temporal signal drift as regressors of no-interest. The *β* weights thus computed were *Z*-scored per voxel, individually for each run. The resulting input data for decoding contained 40 samples (one normalized *β* score per condition per run) for each participant. Separate decoding analyses were conducted for the portions of the V1 ROI in each hemisphere, with an identical model matrix, but using the fitted *β* weights of the regressors corresponding to the stimulation in the contralateral hemifield as input for the classifier.

As in Sengupta et al. (2017), linear support vector machine classifiers (SVM; PyMVPA’s *LinearCSVMC* implementation of the LIBSVM classification algorithm; Chang and Lin, 2011) were used to perform a within-subject leave-one-run-out cross-validation of 4way multi-class orientation classification. Although it has been argued that repeated random splits are a superior validation scheme (Varoquaux et al., 2017), the procedure has been kept identical to Sengupta et al. (2017) in order to facilitate comparability with previously published results. The SVM classifier’s *C* hyper-parameter was, again, tuned using a nested cross-validation approach, where the respective training dataset for each fold of the outer cross-validation iteration was repeatedly split again (leave-another-run-out) to determine an optimal parameter setting by testing 200 equally spaced values from the range [10^−6^, 5] (range enlarged in comparison to the previous study due to search range exhaustion, see Sengupta et al., 2017, Figure S6).

Any results from the previously published 7 Tesla data (Sengupta et al., 2017) that are presented in this study are the result of a re-analysis using the exact same procedure that is described here for the 3 Tesla data.

## 3. Results

### Decoding accuracy and field strength

We computed congruent decoding analyses for the acquired 1.4mm, 2.0 mm (both acquisitions) and 3.0 mm fMRI data for both field strengths. All reported accuracies correspond to the out-of-sample generalization performance of the trained classifiers averaged across participants and hemispheres. Indicators of variation, such as error bars, reflect the between-subject variation of a metric.

Figure 1A shows the comparison of the decoding performance between 7 Tesla and 3 Tesla data using *all* voxels in the individually delineated V1 ROIs as input into the analysis. The decoding accuracies at 7 Tesla exceeded those achieved at 3 Tesla for all resolutions. This difference was observed despite only minimal discrepancies between ROI sizes across field strengths (see Table 1 and Sengupta et al. (2017, Table 1) for comparison). Moreover, there were no systematic differences in the participants’ performance of the attention control task that could explain a performance drop for individual spatial resolutions, or the multiband acquisition (*F*(3,18) = 1.14,*p* = 0.36). Despite this fact we observed notable differences in decoding performance. Peak performance at 3 Tesla was achieved for the 3 mm acquisition. Performance on 1.4 mm data was comparable to the 2 mm multiband acquisition. However, the regular 2 mm acquisition (with GRAPPA) showed near-chance performance when using all V1 voxels as classifier input.

**Figure 1:**
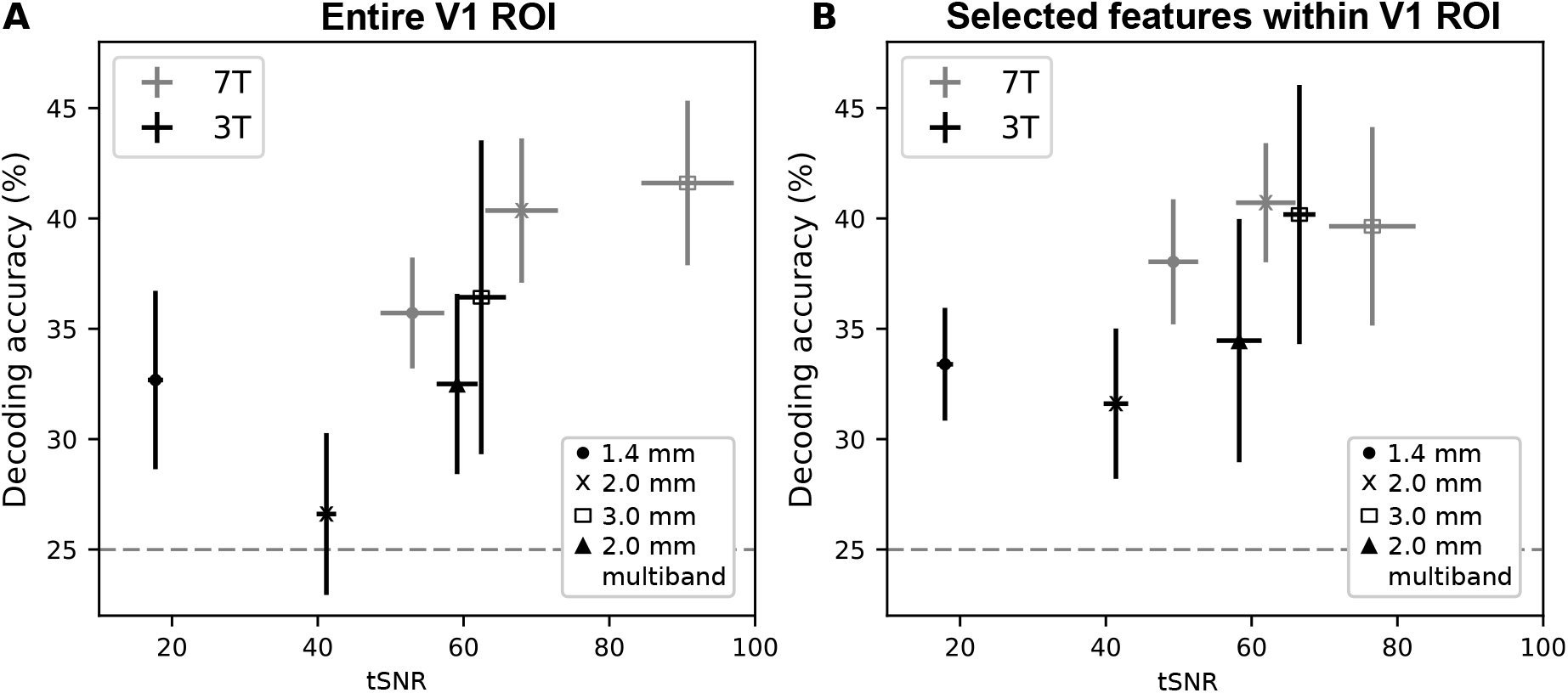
Comparison of orientation decoding performance vs. temporal signal-to-noise-ratio (tSNR) across magnetic field strengths (7 Tesla vs 3 Tesla) and across different acquisition resolutions in the entire retinotopically delineated V1 ROI (A), and after univariate feature selection within V1 (B). Error bars show the SEM for tSNR and accuracy across participants. The maximum decoding accuracy for 3 Tesla data was observed for the 3 mm acquisition. After feature selection peak decoding accuracies for 3 Tesla and 7 Tesla are highly similar, albeit only the 3 mm 3 Tesla analysis yielding a performance similar to the 2 mm or 3 mm 7 Tesla data. The dashed lines indicate the theoretical chance-level.

**Table 1:** V1 ROI size. Average number of voxels for both hemispheres with standard deviation across participants. The effect of univariate feature selection shown as a function of ROI size in 3 Tesla and 7 Tesla datasets.

| Resolution | Delineated V1 (3 Tesla) |  |  |  | After feature selection in V1 |  |  |  |
| --- | --- | --- | --- | --- | --- | --- | --- | --- |
|  | Left hemisphere |  | Right hemisphere |  | 3 Tesla ROI |  | 7 Tesla ROI |  |
|  | #voxels | std | #voxels | std | #voxels | std | #voxels | std |
| 1.4 mm | 1963 | 330 | 1942 | 313 | 901 | 331 | 3044 | 552 |
| 2.0 mm | 831 | 167 | 794 | 140 | 632 | 162 | 1309 | 373 |
| 3.0 mm | 359 | 90 | 347 | 62 | 302 | 67 | 473 | 105 |

In comparison to the report of Sengupta et al. (2017), the larger search range for the *C* parameter optimization resulted in a boost of decoding accuracy for 3 mm 7 Tesla data (results for both other resolutions were replicated), now comparable to the performance on 2 mm data. This has to be considered as evidence against an optimality of 2 mm acquisitions, and attributes the previously reported difference to the particular implementation of the decoding analysis.

To investigate whether the difference in performance can be explained by differences in the available signal between field strengths, we calculated the mean BOLD signal change across all voxels in the V1 ROI in response to each orientation stimulus type across resolutions using FeatQuery in FSL (v5.0.8; Smith et al., 2004). Orientation-related BOLD response magnitude differences are one possible type of signal enabling orientation decoding (Furmanski and Engel, 2000; Tong et al., 2012; Alink et al., 2013). For consistency with the preprocessing for decoding, no spatial smoothing was applied for this analysis. Figure 2A-B shows a substantial difference in the mean V1 BOLD signal magnitude in response to stimulation with oriented flickering gratings between 7 Tesla and 3 Tesla acquisitions. Figure 2A documents a replication of the previous finding of increasing response magnitude with higher resolutions (compare Figure 3B, Sengupta et al., 2017, the global reduction in signal magnitude is explained by the fact that here we report the average response across the entire V1 ROI, while previously only the average of responsive voxels was shown). Figure 2B, however, indicates the absence of any systematic resolution-related magnitude differences in the average ROI signal for 3 Tesla data. A two-factor repeated-measures ANOVA showed neither main effects of acquisition resolution (including the 2 mm multiband acquisition; *F*(3,18) = 0.22,*p* = 0.88), or visual orientation (*F*(1, 6) = 1.453,*p* = 0.27), nor an interaction between the factors (*F*(3,18) = 2.142,*p* = 0.13).

**Figure 2:**
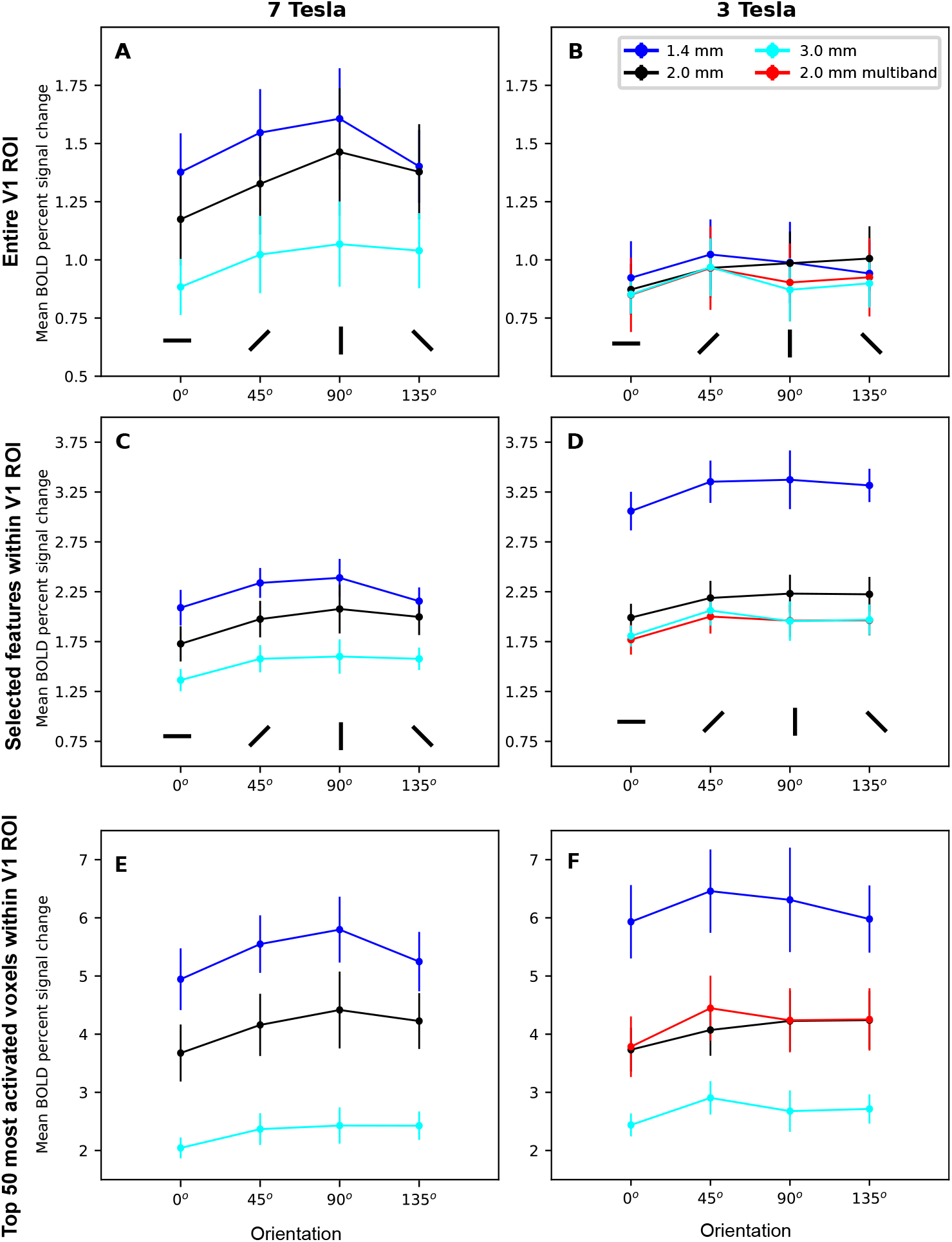
Estimated BOLD signal change by visual orientation for all resolutions at 7 Tesla and 3 Tesla for the retinotopically delineated V1 ROI (A-B) and after univariate feature selection within V1 (C-D). At 7Tesla the average BOLD signal change in V1 increases with higher acquisition resolutions (A). However, this trend was not observed for the average signal change in the entire V1 ROI at 3 Tesla (B). Considering only “responsive” voxels, as determined by univariate feature selection (see main text for procedure), the pattern of increasing signal change observed at 7 Tesla can now be found for both field strengths, at a generally elevated level. In both cases, the signal change observed in the multiband acquisition is similar to that of the 3 mm data. Notably, the average signal change differences for the 1.4 mm vs. 2 mm acquisitions at 3 Tesla are substantially larger than those observed at 7 Tesla. The number of voxels for the comparison between panels *C* and D differs by a factor of 1.5-3 across field strengths (see Table 1). Panel E-F show the average BOLD signal change per orientation in the 50 respective voxels that exhibit the highest average responses to *any* stimulation in the whole V1 ROI, determined for each resolution and field strength separately. The observed average BOLD signal change in those voxels is similar across field in both 3T and 7T data.

**Figure 3:**
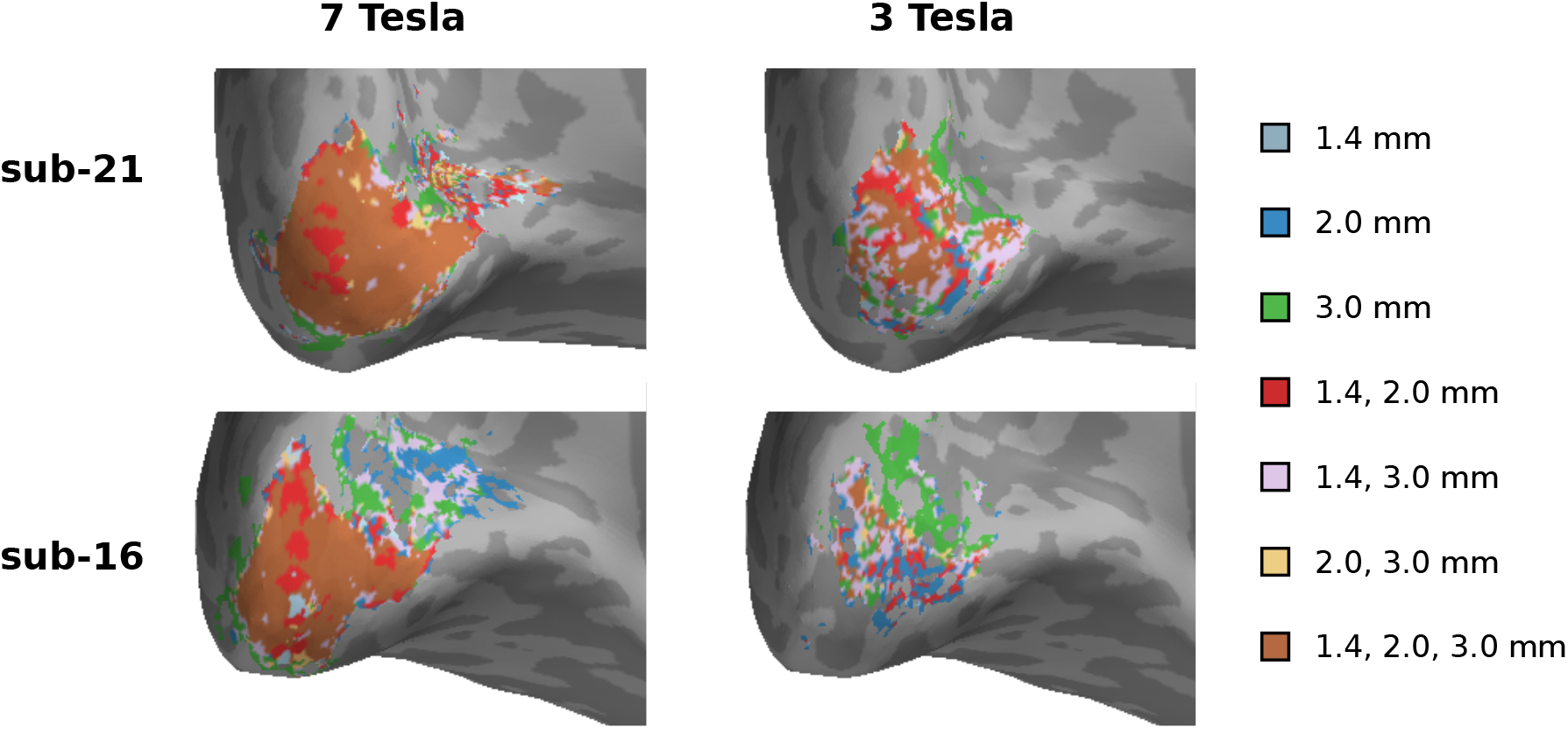
Illustration of the spatial configuration of selected features within the V1 ROI across all resolutions, for both field strengths and two example participants. The color code indicates for which set of resolutions a location was part of the voxel selection. At 7 Tesla, voxel selection patterns appear to be more stable across resolutions, and resulted in a larger number of selected voxels (Table 1). For visual clarity the 2 mm multi-band acquisition is not shown here.

The absence of any visible resolution-related differences could be interpreted as an indication that any available signal is dominated by noise. This would not be entirely surprising, and, in fact, feature reduction/selection, *i.e.*, the exclusion of noisy inputs, is a common component in the application of decoding analysis to fMRI data (Pereira et al., 2009). In order to evaluate this issue, we implemented a univariate feature selection procedure to localize voxels within the V1 ROI that responded to *any* stimulation. A two-level, fixed-effects GLM analysis across all 10 experimental runs was performed for each participant, resolution, and field strength in FSL FEAT (v5.0.8; Smith et al., 2004) (high-pass temporal filtering with a 100 s cutoff, no spatial smoothing, parametric fit of all orientation regressors combined (contrast weights set to 1.0, all others 0.0), cluster threshold *z* > 2.3 with *p* < 0.05). This selection procedure did not involve any comparison or contrast between any combination of orientation stimulation conditions, hence feature selection was performed only once per resolution and field strength, prior to the cross-validated decoding analysis. This setup ensured that all analyses for individual spatial filter configurations are based on the identical set of input voxels to enable valid comparison. Figure 3 illustrates the variability of voxel selections for both field strengths and two exemplar participants. Again, the selection pattern observed for 3 Tesla is visibly noisier compared to 7 Tesla data. The effect of voxel selection on the ROI size is documented in Table 1, leading to substantially smaller ROIs for 3 Tesla – with decreasing differences for increasing resolutions – again indicating a higher noise level.

All data analyses were repeated for both field strengths using this additional feature selection step. Figure 1B shows the decoding performance after feature selection. Mean accuracies for 3 Tesla data increased, in particular for the regular 2 mm acquisition (near-chance performance without feature selection), and for 3mm data. On 3mm data, the performance is now comparable to that on 7 Tesla data. For unfiltered 7 Tesla data, feature selection shows little impact on the decoding performance (in particular for the two lower resolutions), despite a uniform ∼30% reduction in the number of input voxels at each resolution (compare Table 1 with Sengupta et al., 2017, Table 1). This effect was observed despite the expected fact that the average signal change in response to each orientation in the selected voxel subset was higher compared to the average in the entire V1 ROI (Figure 2A vs. C). For 3 Tesla data the effect of voxel selection on BOLD signal magnitude was even more substantial with a clear increase in the average signal of “responsive” voxel with higher resolutions (Figure 2B vs. D), dominated by a boost of the average BOLD signal magnitude for the 1.4 mm acquisition that even exceeds that at 7 Tesla. After voxel selection the structure of signal magnitude difference now resembles those found for 7 Tesla data. The 2 mm multiband acquisition also showed a comparable increase of average signal change in the selected voxel subset, but exhibits the overall lowest signal change at 3 Tesla on the same level as the 3 mm acquisition. Another two-factor (orientation and resolution) repeated-measures ANOVA for the estimated BOLD signal change at 3 Tesla after feature selection revealed a significant effect of acquisition resolution (including the multiband acquisition; *F*(3,18) = 40.73,*p* = 3 · 10^−8^), but again no effect of orientation (*F*(1, 6) = 4.64,*p* = 0.074), or an interaction (*F*(3,18) = 1. 049,*p* = 0.4). However, with this method of analysis, the corresponding number of selected voxels at 3 Tesla is less than a third of those at 7 Tesla, making a direct comparison difficult. For comparison, the average signal change was also computed for the 50 voxels with the largest BOLD response to *any* orientation (Figure 2E and F) for the respective resolution and field strength, revealing a comparable pattern of BOLD signal change magnitudes for both field strengths.

### Effect of temporal signal to noise ratio (tSNR)

According to Chaimow et al. (2011), overall contrast-to-noise ratio (OCNR) impacts decoding performance. The overall contrast-to-noise ratio is inversely proportional to the noise level, where the noise level can be calculated as the inverse of tSNR, which in turn depends on voxel size (Triantafyllou et al., 2005). Previous studies have shown that noise in fMRI time-series is predominantly caused by physiological and thermal noise, and varies primarily with signal intensity, echo time of the EPI sequence and magnetic field strength (Triantafyllou et al., 2005).

Figure 1 documents the relationship of tSNR, field strength, and acquisition resolutions. tSNR magnitudes observed in this study are comparable to those reported by Triantafyllou et al. (2005, Figure 6), indicating a typical noise level for both 3 Tesla and 7Tesla data. Except for the 3 mm acquisitions at both field strengths, the voxel subset determined by feature selection only exhibits marginally different average tSNR compared to the full V1 ROI. For both 3 mm acquisitions the tSNR is higher in the selected voxels than on average in the full V1 ROI.

We fitted a model proposed by Triantafyllou et al. (2005) to characterize the asymptotic relation of tSNR as a function of voxel volume and MR field strength:

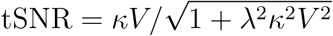

where *V* is the voxel volume, *κ* is the proportionality constant, and λ is the magnetic field strength independent constant parameter. The results for the average tSNR calculated for the full V1 ROI indicate that the data for both field strenghts is compliant with this model (7Tesla: λ=0.013, *κ*=21.94, *R*^2^=0.81; 3 Tesla: λ=0.01, *κ*=6.59, *R*^2^=0.99.

We previously reported that increased tSNR does not necessarily lead to an increase in decoding accuracy (Sengupta et al., 2017). The present results provide further evidence for a more complex or indirect relationship of tSNR and decoding accuracy. At 7 Tesla we did not replicate the finding of a drop in decoding accuracy when lowering the resolution from 2mm to 3mm (see above), however, at the same time we found no evidence for an increase in accuracy, with or without feature selection, despite substantial increase in tSNR for the 3 mm acquisition. At 3 Tesla we observed approximately comparable decoding performance for the 1.4 mm and 2 mm multiband acquisitions, despite an approximately three times larger tSNR for 2 mm data. The same 2 mm multiband acquisition yielded visibly worse decoding performance on the whole V1 ROI than the 3 mm acquisition, despite a similar tSNR level of about 60.

This study included two 2 mm acquisitions, one with and one without parallel imaging acceleration (via GRAPPA) to evaluate the impact of increased tSNR at a constant spatial resolution. Mean tSNR for the non-GRAPPA acquisition was 59.11, compared to 41.17 with GRAPPA factor 2, showing the expected tSNR reduction (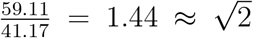). For the decoding analysis on the whole ROI we observed a substantial accuracy gain of more than 10 percentage points, elevating the performance from near-chance level. However, a substantially reduced effect is observed for analysis using the feature selection step, despite practically unchanged tSNR differences between the two acquisitions in the selected voxel set (Figure 1B).

### Spatial scale of the orientation-related signal

In order to get insight into the spatial characteristics of the orientation-related BOLD signal, we applied different types and sizes of spatial Gaussian filters to the data and repeated the identical decoding analysis after each individual pre-processing, analog to Sengupta et al. (2017). LP volumetric Gaussian filtering is commonly applied in fMRI data pre-processing pipelines to reduce high-frequency noise, complementary HP filtered images contain the spatial signal that is removed by the matching LP filter, BP filtered images show how much relevant information is present at a particular spatial frequency band (here with 1mm FWHM bandwidth), and lastly BS filtered images contain the information available outside a particular band. Figure 4 complements Sengupta et al. (2017, Figure 4) with the analog decoding analysis on 7Tesla V1 voxels with and without feature selection using the exact same cross-validation setup as for the 3 Tesla data analyses in this study, with the matching results for all 3 Tesla acquisitions shown in Figure 5.

**Figure 4:**
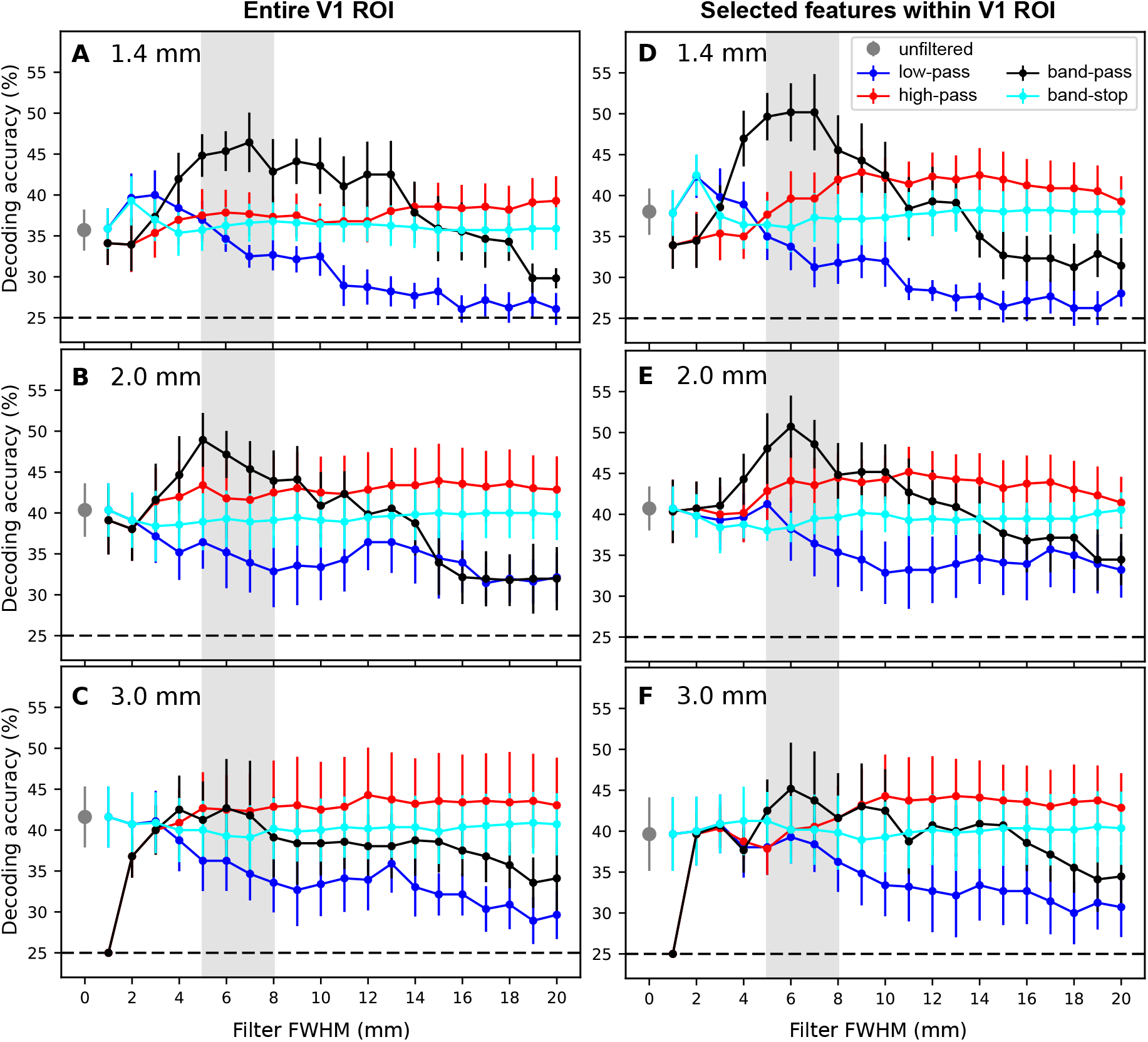
Impact of volumetric Gaussian spatial smoothing on the data acquired at 7 Tesla for the entire V1 ROI (A-C) and the voxel subset determined by univariate feature selection (D-F). The dashed line indicates the theoretical chance level. The patterns of decoding accuracy was similar with and without feature selection across different resolutions. The low-pass components show monotonic decrease in accuracy with increasing in smoothing levels. The high-pass components perform better than the low-pass components beyond 5mm FWHM. The band-pass components showed peak accuracy between 5-8mm FWHM band. The peak in decoding accuracy was more pronounced after feature selection (mostly evident in 1.4mm and 2mm data).

**Figure 5:**
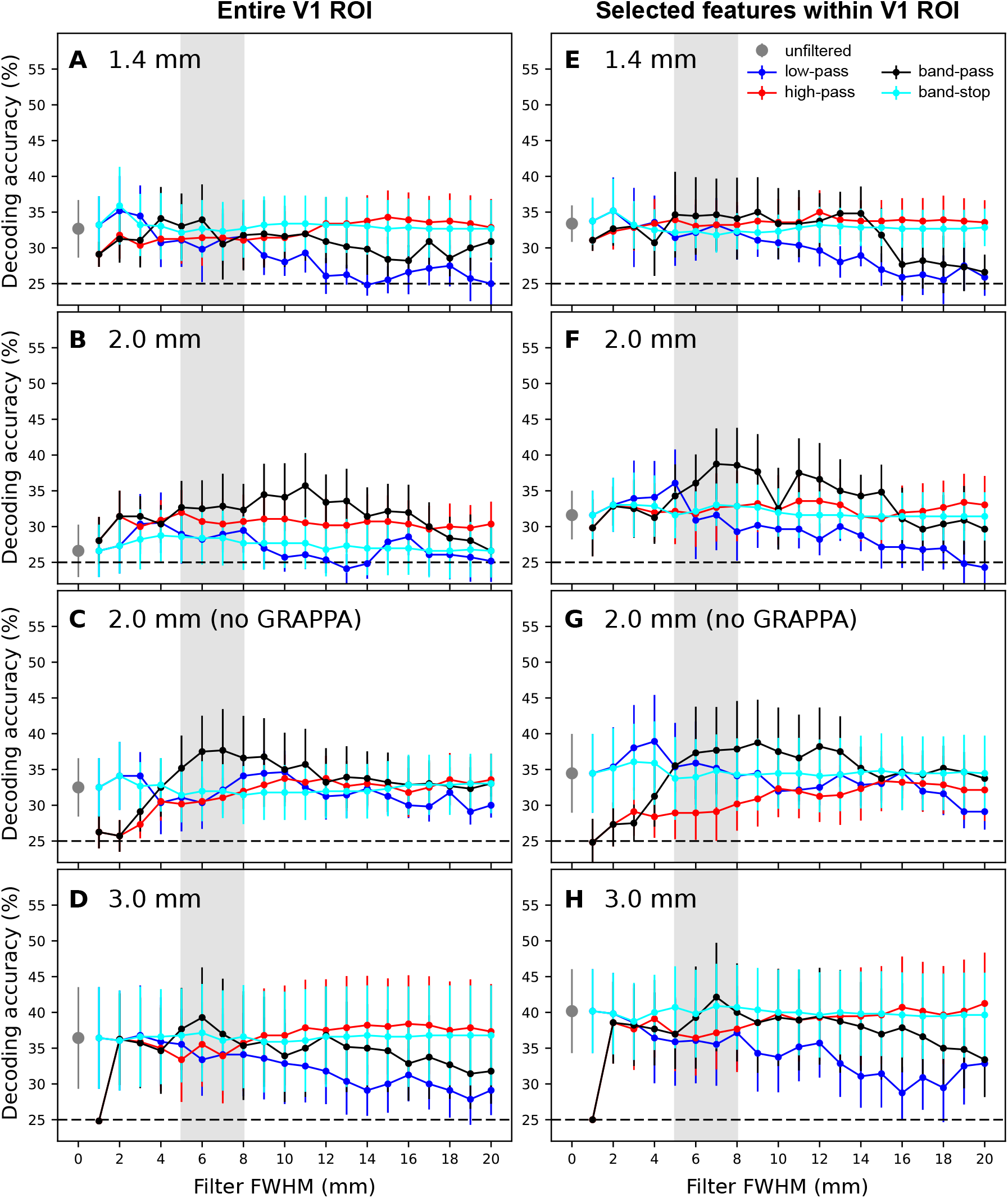
Impact of volumetric Gaussian spatial smoothing on the data acquired at 3 Tesla for the entire V1 ROI (A-D) and the voxel subset determined by univariate feature selection (E-H). The dashed line indicates the theoretical chance level. The variability in the accuracy results across subjects was higher than for the 7 Tesla data, showing the contribution of increased noise in the data. Similar patterns of decoding accuracy across different levels of smoothing were found as in the 7 Tesla data for LP, HP and BS components. The peak in 5-8mm band was not observed for band-pass components for the 1.4mm data. For the 2mm acquisition with and without parallel imaging accelaration, and the 3mm data showed a trend of increased decoding accuracy in the same band but not as pronounced as in the 7 Tesla data.

For all analyses involving the feature selection step, spatial filters were applied first, and voxel time series were selected from the filtered data, based on the pre-computed selection mask for the respective participant and acquisition resolution. Consequently, the selected voxels did not vary across filter types and sizes, and their time series in the filtered data were identical regardless of whether feature selection was applied for a particular decoding analysis.

The pattern of results observed for 7 Tesla data without feature selection replicates our previous report regarding the broad-band nature of the orientation-related signal available for decoding (Sengupta et al., 2017), with peak accuracies being achieved in the ≈5-8 mm band for all tested resolutions, and a detrimental effect of low-pass filtering over high-pass filtering beyond kernel sizes of ≈5 mm. The impact of the change in *C* value search range for classifier tuning was minimal with a tendency towards higher accuracies (compare to Sengupta et al., 2017, Figure 4B-D). Feature selection improves peak decoding accuracies, with increasing impact for higher spatial resolutions (i.e., a larger number of voxels in the ROI prior feature selection), and in particular in the ≈5-8mm band (Figure 5A-C vs. D-F).

The general pattern observed for decoding 3 Tesla data without feature selection resembles that of the 7 Tesla data, albeit at a substantially higher noise level. LP volumetric Gaussian filtering showed a general trend towards a monotonic decrease in decoding accuracy with increase in kernel size, but the decoding performance remained above chance-level even after spatial smoothing of 10 mm, except for the regular 2 mm acquisition without feature selection that already shows a quasi-chance performance without any LP filter applied (Figure 5B). HP components showed above chance decoding accuracy and performed better than the LP filtered image beyond the 9-10 mm filtering kernel in all resolutions, in comparison to the ≈5mm boundary observed for 7 Tesla data. Overall, peak accuracies were obtained on BP filter images.

In comparison to the 7 Tesla data, differences in accuracies between matching resolutions increase substantially in favor of the 7 Tesla data with higher resolutions. This effect was amplified by feature selection. On the other hand, performance on the coarsest tested resolution show little effect of field strength or feature selection. The BP accuracies generally peak in the ≈5-8mm range for all resolutions. This observation is consistent with previous reports for 7Tesla and 3Tesla data (Sengupta et al., 2017; Alink et al., 2013). Classification performance of BS filtered data remained above-chance for all spatial frequency bands, and is approximately equal to the performance in unfiltered data. The BS performance curve initially follows the LP performance for small filter sizes, but resembles the HP performance for larger filter sizes.

## 4. Discussion

The primary focus of this study was to investigate the effect of MR field strength on orientation decoding from primary visual cortex (V1) in order to complement the previous report of Sengupta et al. (2017) on the effect of acquisition resolution at 7Tesla. With that objective, cross-validated orientation decoding analyses using a linear SVM classifier were performed on fMRI data acquired from seven participants at both 7Tesla and 3Tesla for three different acquisition resolutions (1.4mm, 2.0mm and 3.0 mm). With a few exceptions the implementations of stimulation and data analyses were identical to that of the previous study. Two notable exceptions were an enlarged search range for classifier hyper-parameter tuning, based on evidence for a suboptimal configuration in the previous study, and the introduction of an optional feature selection step. In order to enable valid comparisons with the results reported in Sengupta et al. (2017), all relevant analyses were repeated for the 7Tesla data using the exact same configuration as for the 3 Tesla data.

Re-analysis of the 7 Tesla data reproduced the general pattern of results reported in Sengupta et al. (2017), with one exception. A drop in decoding accuracy with a decrease in resolution from 2 mm to 3 mm when using all voxels in the ROI as input was not observed again, and has to be attributed to an inappropriate setup of the classifier hyper-parameter tuning. Therefore the previously proposed optimal tradeoff between acquisition resolution or number of input features and tSNR for 2 mm acquisitions is not supported by the present results. Instead, when viewed in isolation, the decoding accuracies obtained on whole ROI 7 Tesla data suggest a tendency of an increase in tSNR leading to an increase in decoding accuracy, up to an asymptotic limit (Figure 1A). However, when reducing the number of input features via selecting voxels with a significant average response, for each acquisition resolution separately, the observed decoding accuracies are further equalized across resolutions. Accuracy for 1.4 mm data exhibits an increase, while average tSNR for those voxels shows no difference. Feature selection on 2 mm data has little impact, and only for 3 mm there is an indication for drop in accuracy associated with the drop in average tSNR for the selected voxels (Figure 1B). Overall, this indicates that the differences in average tSNR observed for 7 Tesla data across resolutions and voxel sets do not explain the observed decoding accuracies. The peak decoding accuracies in the ≈5-8mm band document that comparable decoding performance can be achieved for 1.4 mm and 2 mm data, regardless of their tSNR differences (Figure 5A-B). This is particularly evident for decoding on the selected voxel set that reduces the discrepancy in the number of input voxels between resolutions (Figure 5D-E).

At 3 Tesla, the two 2 mm data acquisitions enable a direct evaluation of the impact of tSNR on decoding accuracy. The manipulation of acquisition parameters by replacing parallel imaging (GRAPPA) with simultaneous multi-slice acquisition yielded the expected increase in tSNR, while maintaining the same field-of-view, resolution, and repetition time across both scans. The results, however, again suggest no strong direct coupling between tSNR and decoding accuracy. While for decoding on the entire V1 ROI accuracy increased substantially (from quasi-chance) with the increase in tSNR for the multiband acquisition, this difference was visibly reduced for decoding on the selected voxel set – despite practically identical tSNR differences in both cases (Figure 1). At the same time, minimal tSNR differences between the 2 mm multiband acquisition and the 3 mm scans translate into a boost of decoding accuracy in favor of the lower-resolution scan. Moreover, decoding performance for the 2 mm multiband scan and the 1.4 mm acquition was comparable, despite a factor 3 difference in tSNR, with or without feature selection (Figure 1). Taken together, these results are congruent with Tong et al. (2012) in indicating that increase in tSNR generally benefits decoding, but the relationship is non-linear, and depends on other factors, such as acquisition resolution, and magnetic field strength.

### BOLD signal change

As reported in Sengupta et al. (2017) for 7Tesla, we again found no statistically significant orientation-specific differences in the average signal change across voxel, in particular not higher signal change for cardinal vs. oblique orientations (see Furmanski and Engel, 2000; Swisher et al., 2010, for reporting of such differences with opposing signs).

Although there was a significant main effect of acquisition resolution on the percentage signal change calculated from the whole V1 ROI in 7 Tesla, this was not observed in the corresponding 3 Tesla data, as expected due to an elevated noise level. To address this issue, a widely used conventional feature selection method was used which favors voxels that exhibit a larger average response to any orientation, and not necessarily voxels that are exhibiting maximum signal difference between a given pair of orientations. This is a possible explanation for the fact that the higher average signal change of the selected voxel set at 3 Tesla did not translate to an increase in decoding accuracy. Consequently, the resulting decoding models might differ substantially regarding the nature of information used to discriminate orientations, despite comparable performance.

As with tSNR, the observed signal change differences across resolutions at both 7 Tesla and 3 Tesla do not explain the associate decoding accuracies in a straightforward manner. For 7T and 3T data with the feature selection procedure, the signal change increases with resolution, but higher signal change does not translate into improved decoding accuracies. These results could be attributed to the fact that signal change was computed as an average across all voxels (Demetriou et al., 2016), and the number of voxels varied substantially across resolutions and field strengths (Table 1). However, when controlling for these differences, by considering only an equal number of voxels with the highest average BOLD response, the signal change pattern observed at 3 Tesla resembles that at 7 Tesla with an increase in signal change with increasing resolution, in contrast to the observed pattern of decoding accuracies. We did not observe a generally larger BOLD signal change at 7 Tesla compared to 3 Tesla. The voxel set determined by feature selection for the 1.4 mm acquisition even exhibits a larger average BOLD signal change at 3 Tesla. This could be, at least in part, a consequence of our particular choice of echo times: TE=22ms (7T) and TE=30ms (3T). BOLD signal change and echo time are linearly related, with larger echo times being associated with larger signal change (van der Zwaag et al., 2009, Figure 4). While this difference would shrink an expected BOLD signal advantage in favor of 3 Tesla the effect of voxel selection is likely to play a comparatively bigger role. Contribution of BOLD signal with venous origin is expected to be proportionally increased at 3 Tesla (van der Zwaag et al., 2009), and case-by-case visual inspection (data not shown) indicates that the selected top 50 voxels for the 1.4 mm acquisition at 3 Tesla are indeed predominantly located within the set of voxels classified as “venous” in Sengupta et al. (2017).

### Spatial scale of orientation-related signal

Similar to our previous report for 7 Tesla (Sengupta et al., 2017) we find that the orientation-related signal at 3Tesla is not confined to a particular spatial frequency band. Increasing LP filter kernel sizes are associated with a gradual decline in decoding accuracy but performance was above chance beyond 10 mm smoothing – evidence for a contribution of coarse scale signals. On the other hand, the HP filtered components beyond 10 mm show a decoding performance comparable to that on unfiltered data, indicating the availability of relevant high frequency signals. The band-stop components always perform above chance level with decoding accuracies similar to the low-pass filtered components until the p¿5-8 mm band, and then resembling the accuracies of the high-pass filtered component. This documents that low frequency and high frequency components can contribute to decoding. Overall, it can be concluded that, at 3 Tesla, the orientation related BOLD signal is spatially broadband in nature, starting from millimeter range columnar signals to large-scale orientation biases (beyond 10 mm). While this general finding is in line with reports for 7Tesla data (Sengupta et al., 2017; Swisher et al., 2010), the direct comparison between field strengths in this study indicates that the spatial characteristics of BOLD signal patterns informative for decoding are shifted towards more coarse-scale patterns at 3 Tesla (compare the crossover point of HP and LP decoding accuracies: 3-5mm (7T) vs. >8mm 3T;Figure 4 and 5). This difference is consistent with the increase in BOLD PSF size for 3Tesla compared to 7Tesla (Shmuel et al., 2007; Engel et al., 1997; Chaimow et al., 2018).

### Practical conclusions

This study was motivated by the observation in Sengupta et al. (2017) that for 7Tesla data “maximum” decoding performance can be achieved at a resolution that is routinely accessible with more conventional 3 Tesla scanners. However, the results reported here indicate that, despite identical spatial acquisition resolutions, data from these field strength differ substantially – within the constraints set by the analysis procedure chosen here. First and foremost, 3 Tesla data are evidently more noisy with respect to orientation decoding. This is reflected in the overall lower decoding accuracies, regardless of resolution. The elevated noise level also translates into a larger variability of voxel selection procedures, which may in turn impair comparability of results across field strengths and resolutions in terms of the particular nature of the signal that is available for decoding. However, this difference in noise at the level of distributed BOLD signal patterns is not immediately evident in global quality indicators, such as average tSNR, or average BOLD signal change magnitude (Formisano and Kriegeskorte, 2012). These empirical results are largely congruent with the conclusions of a simulation study by Chaimow et al. (2018) that attributes the finding of superior decoding performance (pattern information) for high-resolution 7T scans primarily to the smaller PSF width. In contrast to those simulation results we did not observe 3 Tesla maximum decoding accuracy at resolutions higher than the lowest (3 mm) resolution tested here. However, a proper comparison to this simulation is difficult, as the true scale and nature of the signal are unknown.

The impact of the adjusted search range for hyper-parameter optimization in this study demonstrates the importance of comparatively “minor” details of a decoding analysis setup. Arguably, many studies that utilized the popular cross-validated SVM decoding performed no hyper-parameter optimization at all (frequently the default *C* = 1 is left unchanged and untested). In Sengupta et al. (2017) a too constrained optimization strategy still led to an interpretation that is now challenged by the reanalysis presented in this report. Clearly, we cannot claim or substantiate in any way that the present analysis strategy is optimal now – but the adjusted procedure used in this study has been applied in an identical fashion to both datasets, so that any results presented here are comparable across field strengths. However, future studies should have a stronger focus on exploring potential confounds of analysis parameters with variables of interest.

Lastly, in this study we replicated the finding from Sengupta et al. (2017) that spatial LP filtering (smoothing), a standard component of many fMRI data analysis pipelines, is not beneficial for decoding visual orientation from V1. Due to the broadband nature of the orientation-related signal, superior decoding performance can be achieved via BP filtering strategies that remove high *and* low frequency noise components. The present results offer no basis for conclusive recommendations on the specific configuration of a BP filter, however, we recently showed that this finding is not limited to orientation decoding from visual cortex, but a similar effect can be observed for decoding musical genre from auditory cortex (Sengupta et al., 2018). Given the potential generality of this finding future studies focused on the analysis of distributed activity patterns using, for example, decoding analysis or encoding models should investigate this possibility.

## Acknowledgements

This research enabled by a grant from the German Research Foundation (DFG) awarded to S. Pollmann and O. Speck (DFG PO 548/15-1). This study was, in part, also supported by the German Federal Ministry of Education and Research (BMBF) as part of a US-German collaboration in computational neuroscience (CRCNS; awarded to J. V. Haxby, P. Ramadge, and M. Hanke), co-funded by the BMBF and the US National Science Foundation (BMBF 01GQ1112; NSF 1129855). M. Hanke was supported by funds from the German federal state of Saxony-Anhalt and the European Regional Development Fund (ERDF), project: Center for Behavioral Brain Sciences.

